# Electrodiffusion analysis of concentration and voltage changes in thin cylindrical domains using cross-diffusion modelling

**DOI:** 10.64898/2026.05.13.724841

**Authors:** Frédéric Paquin-Lefebvre, Jürgen Reingruber

## Abstract

A major challenge in neuroscience is to predict how currents in nanodomains affect voltage and ionic concentrations. Cable and Rall theory provide analytic current-voltage relations by neglecting concentration gradients, and the impact of concentration gradients is usually studied numerically with the Poisson–Nernst–Planck (PNP) model. A precise quantitative understanding of the combined dynamics remains limited because analytic current-voltage-concentration relations are missing. In this work we derive such relations using a novel approach based on cross-diffusion equations. For narrow cylindrical domains, we derive time-dependent and steady-state expressions that explicitly show how currents affect voltage and ionic concentrations. We find that the influx of only one ion can significantly change the concentrations of all the other ions even if no channels for these ions are present. After a current injection we compute a biphasic voltage transient where the small-time asymptotic corresponds to the steady-state solution of the cable equation. We show that the accuracy of cable theory prediction for the voltage depends on how the current is distributed among the various ions. Finally, we develop an iterative method to accurately compute steady-state profiles for voltage and concentrations using first-order results by subdividing a cylinder into small segments.

## 1 Introduction

Understanding the electrical activity in neurons is key to comprehend the brain functioning. Steady improvements in experimental techniques allow to quantify neuronal activity with nanoscale resolution [1, 2]. For example, the invention of the voltage-clamp technique [3, 4] led to the discovery of the ionic basis of action potentials [5]. Recent advances in genetically encoded voltage indicators [6, 7] and nanopipette technology [8, 9] reveal spatiotemporal voltage patterns in neuronal nanodomains. Contrary to electrical circuits that depend only on current-voltage relations, the electrical response of neurons also depends on the dynamics of ion concentrations [10, 11]. But measuring ion concentrations in small domains is much more challenging compared to voltage. Since many key functions in neurons are controlled by *Ca*^2+^, advanced methods have been developed to measure the intracellular *Ca*^2+^-concentration with chemical fluorescent indicators or genetically encoded calcium indicators [12, 13]. The *Na*^+^ and *K*^+^ concentration dynamics is often neglected and reduced to generating driving forces for currents between intracellular and extracellular spaces, for example in the famous Hodgkin-Huxley model for action potential generation [5]. Recent experimental results, however, show that the *Na*^+^ and *K*^+^ dynamics matter for the proper functioning of neuron and glial cells [14, 15, 16, 17]. In tiny domains even small currents can significantly alter ionic concentrations and voltage. For example, a 10*pA Na*^+^-current flowing into 1µm^3^ volume corresponds to the flux *J* = *I/*(*ℱ/V*) ∼ 100*mM*/*s* (with the Faraday constant *ℱ* ≈ 0.1*spA*/(*mMµm*^3^)), which has the potential to change the *Na*^+^ concentration by 100*mM* in one second. The most accurate electrodiffusion models are based on Nernst-Planck expressions for ionic fluxes and the Poisson equation for the voltage [11]. In neuroscience, the Poisson-Nernst-Planck (PNP) model has been applied to study transport through ion channels [18, 19, 20, 21], Debye-layer properties near membranes [22, 23, 24], the propagation of action potentials [25, 26] or electrical properties in small mitochondrial cristae, cardiac dyads and dendritic spines [27, 28, 29, 2, 30, 31] (see [32] for a recent review). Because PNP models are difficult to study even numerically, more coarse grained approaches have been developed. Charge-capacitor models maintain the Nernst-Planck expressions for ionic fluxes but replace the Poisson equation by an effective equation where the potential change is determined by the change of the total charge and the membrane capacitance [33, 34, 35, 36, 37, 38]. Finally, the cable equation and the Rall theory of a neuron are based on constant ion concentrations, in which case electrical properties are determined by the coupling between voltage and currents [39, 40, 41, 42]. Rall theory and capacitance based voltage models that include reaction–diffusion equations are the core of popular simulation software such as NEURON [43] or ASTRO [44].

In electrical models of neurons membrane currents flowing into a volume usually directly alter only the concentrations of ions that contribute to these currents. For example a *Na*^+^ current changes only the *Na*^+^ concentration, and might indirectly affect other ions by modifying voltage or concentration dependent channel properties. However, this approach ignores that the concentrations of all ions are coupled via the Poisson equation and electroneutrality constraints. So far, a precise quantitative understanding of the implications of this coupling is lacking because PNP models are usually studied only numerically due to their complexity. In this work we provide a quantitative understanding by deriving analytic results that explicitly show how axial and membrane currents affect ion concentrations in narrow cylindrical domains. We use the electroneutrality approximation to eliminate the Poisson equation, and we derive a system of cross-diffusion equations that expose how the concentration of a specific ion is affected by currents and concentration gradients of other ions. We explicitly solve these equations for constant current injections and in the limit of small concentration changes, and we derive first-order solutions for transient and steady-state profiles of concentrations and voltage. We find that the steady-state cable solution describes the small-time asymptotic of the voltage. We further derive higher-order steady-state results, and we devise a recursive method to accurately compute steady-state profiles based on first-order results by subdividing the cylinder into small segments. Finally, we investigate several biological scenario with different current injections, and we verify our analytic results with numerical simulations of the exact 1D PNP model.

## 2 Electrodiffusion model

We consider electrodiffusion in a spatially uniform cylindrical domain of radius *R* and length *L* with volume *V* = *πR*^2^*L* and surface *S* = 2*πRL*. We normalize distances by *L* such that the longitudinal position along the cylinder is given by the normalized variable 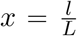, with 0 ≤ *x* ≤ 1. We assume rotational symmetry and fast radial equilibration such that quantities depend only on the time t and the longitudinal position *x*. We consider the normalized voltage 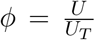, where 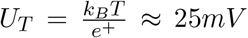. We include various species of mobile ions labelled by the index *s* and characterized by the valences *z*_*s*_, diffusion constants *D*_*s*_, and concentrations *c*_*s*_(*x, t*) or charge densities *ρ*_*s*_(*x, t*) = *ℱV z*_*s*_*c*_*s*_(*x, t*). We further have a uniform background charge density *ρ*_*b*_ due to immobile ions. Concentrations change due to longitudinal fluxes *J*_*s*_(*x, t*) and membrane fluxes *J*_*m,s*_(*x, t*). The membrane and axial currents associated with these fluxes are *I*_*m,s*_(*x, t*) = *z*_*s*_*ℱJ*_*m,s*_(*x, t*)*S* and *I*_*s*_(*x, t*) = *z*_*s*_*ℱJ*_*s*_(*x, t*)*πR*^2^. We use the convention that inward fluxes are negative and outward fluxes positive. Thus, inward fluxes of cations and outward fluxes of anions generate depolarizing negative currents. With the Nernst-Planck expression 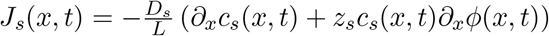 [11] we obtain the axial currents

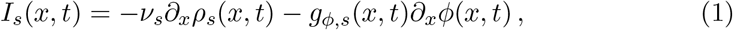

where *v* = *D*_*s*_ /*L*^2^ are diffusion rates and 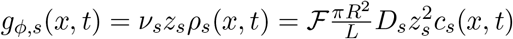 are cytoplasmic conductivities. Eq. 1 is a non-ohmic relation between current and potential due to concentration gradients. The conservation equations for the charges are

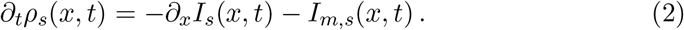

To complete the model we further need an equation for the potential. The classical choice is the Poisson equation [45]

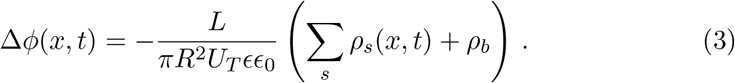

Eqs. 2 and 3 are known as Poisson-Nernst-Planck (PNP) equations. We numerically solve the PNP equations with COMSOL Multiphysics, but to derive analytic results we replace the Poisson equation with the electroneutrality condition

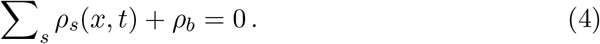

Eq. 4 is motivated by Eq. 3 because 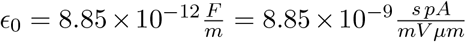 is small. An alternative approximation consists in replacing the Poisson equation with the charge-capacitor equation *SC*_*m*_*U*_*T*_ *∂*_*t*_*ϕ*(*x, t*) = −*∂*_*x*_*I*(*x, t*) −*I*_*m*_(*x, t*), with membrane capacity *C*_*m*_ = 1*µF*/*cm*^2^ = 10^*−*5^*spA*/(*mV µm*^*2*^) [33]. The charge-capacitor equation for the voltage is formally identical to the cable equation, except that currents also depend on concentration gradients. For *C*_*m*_ → 0 we recover the electroneutral model [36], which has been found to provide very good approximations to the PNP model [46, 36].

## 3 Time dependent analysis

Next we use the electroneutral model to investigate how initially uniform distributions of potential and concentrations inside a cylinder change as a function of membrane currents *I*_*m,s*_(*x, t*) and tip currents *I*_*tip,s*_(*t*) injected for *t* > 0 (Fig. 1). At *x* = 0 the cylinder is connected to a large reservoir (e.g cell body), and here we therefore assume Dirichlet boundary conditions *ϕ* = 0 and *c*_*s*,0_. At the tip *x* = 1 we have a reflecting boundary with injected currents.

**Figure 1:**
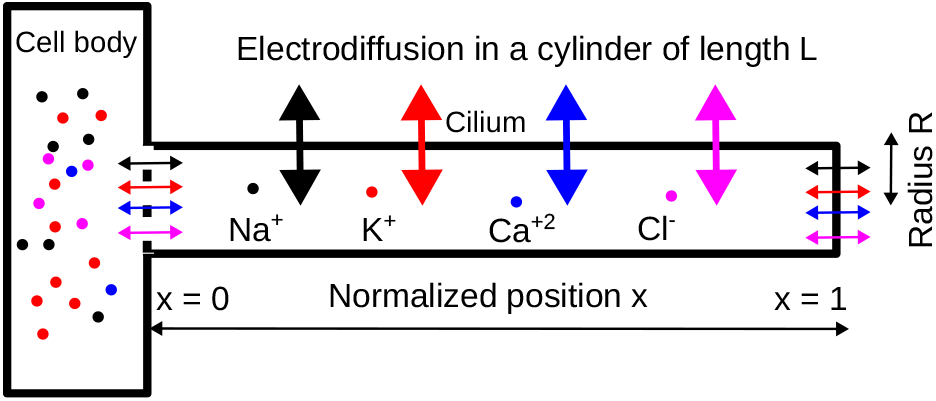
Schematic of the electrodiffusion model. A thin cylinder (e.g. dendrite or cilium) is connected to a large reservoir (e.g. cell body) where voltage and concentrations are constant. For applications we use *R* = 0.1*µm* and *L* = 25*µm*.

By integrating Eq. 2 using the electroneutrality condition *∂*_*t*_ *∑*_*s*_ *ρ*_*s*_(*x, t*) = 0 we obtain the total axial current

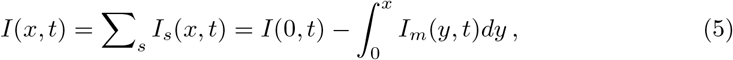

with 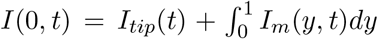, *I*_*tip,s*_(*t*) = ∑_*s*_ *I*_*tip,s*_(*t*) and *I*_*m*_(*y, t*) =∑_*s*_ *I*_*m,s*_(*y, t*). Whereas *I*(*x, t*) is determined by the injected currents, this is not the case for *I*_*s*_(*x, t*) because *∂*_*t*_*ρ*_*s*_(x, t)≠ 0. From Eq. 1 we get *g*_*ϕ*_(*x, t)∂*_*x*_*ϕ*(*x, t*) = −*I*(*x, t*) −∑_*s*_ *v*_*s*_*∂*_*x*_*ρ*_*s*_(*x, t*) (*g*_*ϕ*_(*x, t*) = ∑_*s*_ *g*_*ϕ,s*_(*x, t*)) and

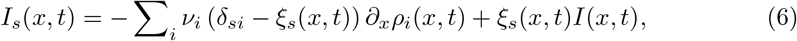

with the Kronecker *δ*_*si*_ and the conductivity fractions

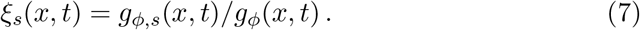

Finally, by combining Eq. 6 and Eq. 2 we obtain for ρ_*s*_(*x, t*) the closed system of non-linear equations

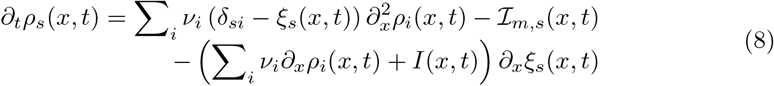

with

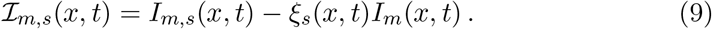

At *x* = 0 we have Dirichlet boundary conditions. From Eq. 6 we find at *x* = 1 the boundary condition ∑ _*i*_ *v*_*i*_ (*δ*_*si*_ − *ξ*_*s*_(1, *t*)) *∂*_*x*_*ρ*_*i*_(*x, t*)|_*x*=1_ = −*ℐ*_*tip,s*_(*t*) with

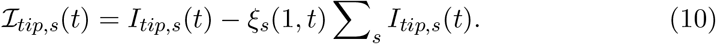

Because Eq. 8 is difficult to solve in general, we derive asymptotic results with a linear first-order approximation of Eq. 8 valid for small concentration changes.

### 3.1 First-order results with small concentration changes

To simplify Eq. 8 we use the first-order approximation

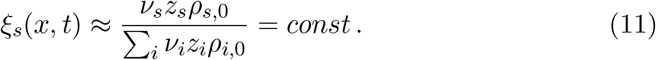

Eq. 11 is valid for small concentration changes inside the cylinder. With Eq. 11 we have that Eq. 8 reduces to the system of cross-diffusion equations

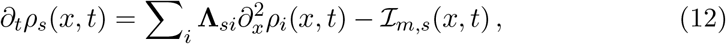

with the matrix **Λ**_*si*_ = *v*_*s*_*δ*_*si*_ − *ξ*_*s*_*v*_*i*_. Eq. 12 links the concentration change of ion s to gradients and currents of other ions. The impact of other ions increases with *ξ*_*s*_. The boundary condition at *x* = 1 is ∑ _*i*_ **Λ**_*si*_*∂*_*x*_*ρ*_*i*_(*x, t*)|_*x*=1_ =−ℐ_*tip,s*_(*t*). **Λ** is singular because 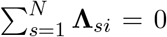. The eigenvectors 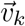, *k* =1, …, *N* − 1, to the *N* − 1 non-zero eigenvalues satisfy ∑_*s*_ *v*_*k,s*_ = 0 (expres-sions for *v*_*k,s*_ are given in Eq. 26 in the appendix). Because ∑_*s*_ ℐ_*tip,s*_(*x, t*) = ∑_*s*_ ℐ_*m,s*_(*x, t*) = 0 we obtain the decompositions

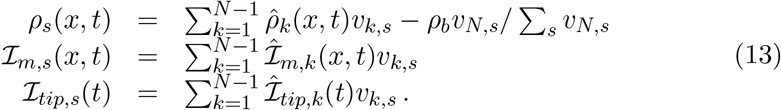

The functions 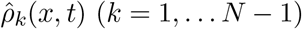 (*k* = 1, … *N* − 1) satisfy

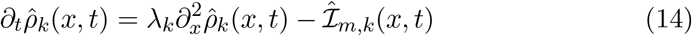

with boundary condition 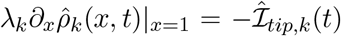. The first-order approximation of Eq. 1 is *g*_*ϕ*_*∂*_*x*_*ϕ*(*x, t*) = −*I*(*x, t*)−∑ _*s*_ *v*_*s*_*∂*_*x*_*ρ*_*s*_(*x, t*) with constant conductivity *g*_*ϕ*_ computed with the concentrations at *x* = 0. We obtain

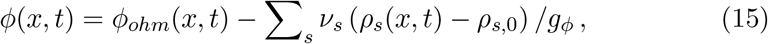

where *ϕ*_*ohm*_(*x, t*) is solution of −*g*_*ϕ*_*∂*_*x*_*ϕ*(*x, t*) = *I*(*x, t*). As a special case, if all ions have the same diffusivity (*v*_*s*_ = *v*) the contributions of concentration gradients cancel and *ϕ*(*x, t*) = *ϕ*_*ohm*_(*x, t*).

Next we solve Eq. 14 for *t* > 0 with constant currents *I*_*tip,s*_ and *I*_*m,s*_. We get

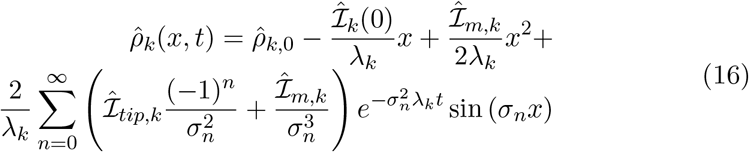

with 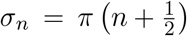 and 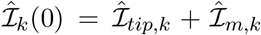. The ohmic potential in Eq. 15 is solution of the steady-state cable equation *I*(*x*) = −*g*_*ϕ*_*ϕ*^*′*^(*x*) with *I*(*x*) = *I*(0) − *I*_*m*_*x*,

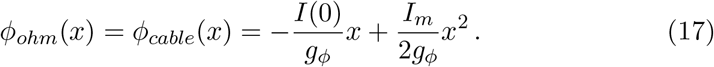

With Eq. 6 we obtain the currents

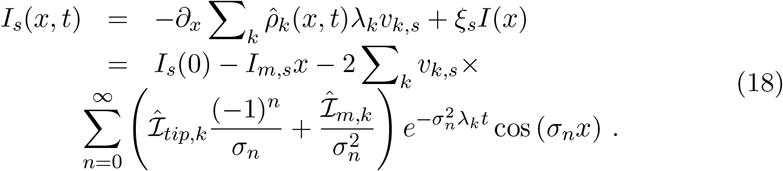

#### Ambipolar diffusion

For two ions with valences *z*_1_ and *z*_2_ the electroneutrality condition implies

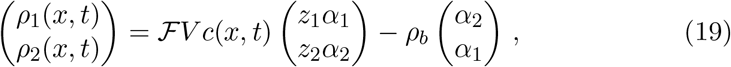

where *c*(*x, t*) = *c*_1_(*x, t*) + *c*_2_(*x, t*), 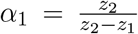 and 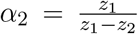. Because (*z*_1_*α*_1_, *z*_2_*α*_2_)^Τ^ is eigenvector of **Λ** with eigenvalue λ_1_ = *ξ*_2_*v*_1_ + *ξ*_1_*v*_2_ (see Eqs. 26-27), Eq. 14 reads

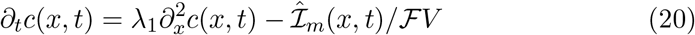

with 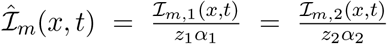. Moreover, with *ρ*_*b*_ = 0 we have 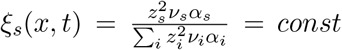, in which case Eq. 11 and Eq. 20 are exact. Eq. 20 without membrane currents is known as ambipolar diffusion equation or Nernst-Hartley equation [47, 48, 49].

## 4 Steady-state results beyond first-order approximation

To go beyond the first order results in Eq. 16 and Eq. 15, we consider Eq. 1 with electroneutrality and obtain for the potential the equation

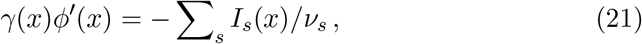

where 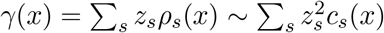is the total ionic strength of the electrolyte. Eq. 21 shows that *ϕ*(*x*) and ∑ _*s*_ *I*_*s*_(*x*)/*v*_*s*_ always satisfy an ohmic relation, whereas *ϕ*(*x*) and *I*(*x*) =∑ _*s*_ *I*_*s*_(*x*) have an ohmic relation if all ions have the same diffusivity *v*_*s*_ = *v*. In this case currents due to concentration gradients annihilate. Note that ϕ(x) and I_*s*_(x) never satisfy an ohmic relation in presence of concentration gradients (Eq. 1).

To obtain an analytic formula for ϕ(x) we solve Eq. 21 with *I*_*s*_(*x*) = *I*_*s*_(0)−*I*_*m,s*_*x* and the quadratic expansions 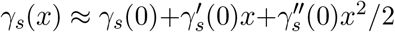. γ_*s*_(0) are given by the Dirichlet boundary condition, and 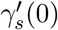 and 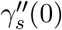 are computed with Eq. 30 in the appendix. The solution of Eq. 21 with γ(*x)* = ∑ _*s*_ γ_*s*_(*x*) = γ_0_(*x* − γ_1_)(*x* − γ_2_) is

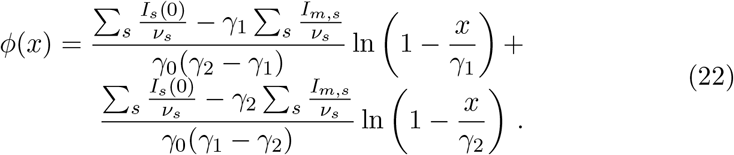

Eq. 22 is the exact for the ambipolar case with *ρ*_*b*_ = 0. With Eq. 1 we compute the concentrations with

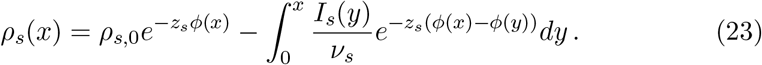

### 4.1 Steady-state results with small segments

Because the first-order results in Eq. 16 and Eq. 15 scale ∼ *L*/*R*^2^, the error increases with the length of a cylinder. To obtain more accurate results we use a recursive method based on first-order results and splitting of the cylinder into *N*_*seg*_ equal segments labelled by 1 ≤ *i* ≤ *N*_*seg*_. Concentrations and voltage in segment i with segmental position 0 ≤ *y* ≤ 1 are

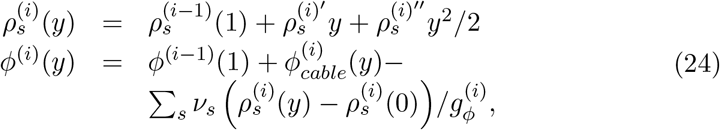

with 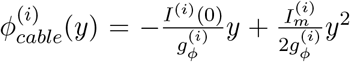. The segmental membrane currents are 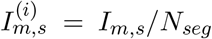. The axial currents flowing into segment i located at cylindrical position 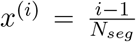 are 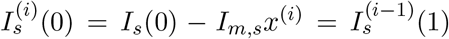. The parameters 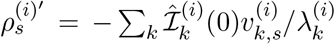and 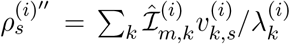 are computed with segment specific matrices **Λ**^(*i*)^. By construction, the overall solution obtained by concatenating the segmental parts are continuous, however, derivatives at segment interfaces are discontinuous because the conductances 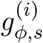 change discontinuously. But these discontinuities can be made very small by reducing the segment length.

### 4.2 Cable equation and constant concentrations

The cable equation assumes that concentrations remain constant. This condition is achieved with electrodiffusion if currents satisfy (Eq. 1 with 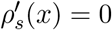)

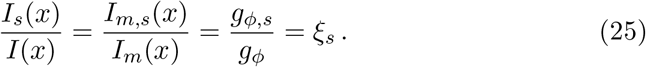

Eq. 21 and Eq. 25 give the steady-state cable equation *I*(*x*) = −*g*_*ϕ*_*ϕ*^*′*^(*x*). Hence, we conclude that using the cable equation to compute *ϕ*(*x*) as a function of *I*(*x*) implicitly implies that *I*_*s*_(*x*) are distributed according to Eq. 25. However, this condition is usually violated, for example, by assuming that *I*(*x*) is a *Na*^+^ current.

## 5 Applications

We now apply our analytic results to investigate how currents affect ionic concentrations and potential in a thin cylindrical domain with radius *R* =0.1 *µm* and length *L* = 25 *µm* (Fig. 1). We consider the dynamics of *Na*^+^, *K*^+^, *Ca*^*2*+^ and *Cl*^−^ with parameters from Table 1. To obtain electroneutrality we also have a uniform background charge density. We compare our analytic results with numerical simulations obtained by solving the PNP equations (2-3) with COMSOL. We solve the 1D Poisson equation Eq. 3 with *ϵ* = 80 corresponding to water. The boundary condition at *x* = 1 is obtained from Eq. 1 with electroneutrality, *ϕ*^*′*^(1, *t*) ∑ _*s*_ *z*_*s*_*ρ*_*s*_(1, *t*) = −∑ _*s*_ *I*_*tip,s*_(*t*)/*v*_*s*_.This boundary condition is consistent with Eq. 21.

**Table 1:**
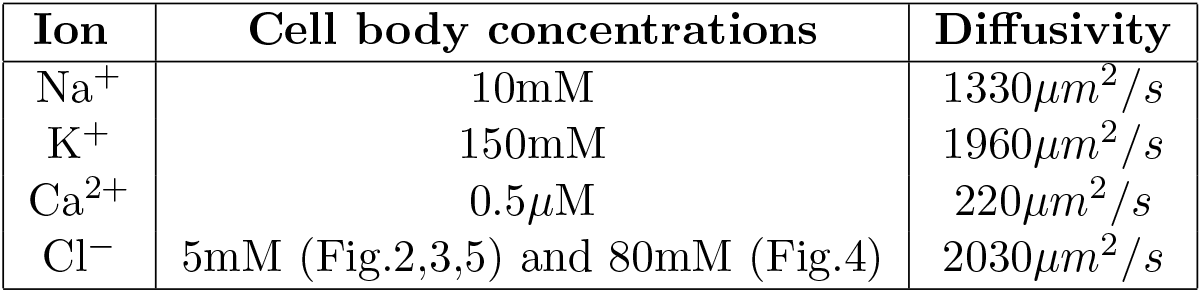
Concentration and diffusivity values [11].

| Ion | Cell body concentrations | Diffusivity |
| --- | --- | --- |
| Na<sup>+</sup> | 10mM | 1330$\mu\text{m}^2/\text{s}$ |
| K<sup>+</sup> | 150mM | 1960$\mu\text{m}^2/\text{s}$ |
| Ca<sup>2+</sup> | 0.5$\mu\text{M}$ | 220$\mu\text{m}^2/\text{s}$ |
| Cl<sup>-</sup> | 5mM (Fig.2,3,5) and 80mM (Fig.4) | 2030$\mu\text{m}^2/\text{s}$ |

We first study the transient dynamics following the injection of constant *Na*^+^ tip and membrane currents for *t* > 0, *I*_*m,Na*_ = *I*_*tip,Na*_ = −5*pA* (Fig. 2). As a general conclusion, with small currents our first-order results in Eq. 16 and Eq. 15 are in very good agreement with PNP solutions (Fig. 2, bullets vs solid, dashed and dotted lines). After switching on the currents at *t* = 0, the voltage jumps from *ϕ*(*x*) = 0 to the steady-state cable equation solution *ϕ*_*cable*_(*x*) given in Eq. 17 (Fig. 2A). This initial voltage transition is instantaneous for the first-order solution *ϕ*(*x, t*) = *ϕ*_*cable*_(*x*) − Σ_*s*_ *v*_*s*_ (ρ_*s*_(x, t) − ρ_*s*,0_) /g_*ϕ*_ (Eq. 15). This can be understood by noting that the electroneutrality condition can be derived from the charge-capacitor approximation in the limit of vanishing membrane capacitance *C*_*m*_ → 0 [36], which entails instantaneous voltage adjustments. The initial voltage change is not immediate but extremely fast for the PNP solution because the permittivity ϵ_0_ is so small (Fig. 2A), which has also been observe in more complex PNP models [30]. Because concentrations remain almost constant during this fast transient, the steady-state cable solution corresponds to voltage profile after the jump (Fig. 2A). After this fast initial adjustment, the voltage is slaved by the concentrations and co-evolves with them in a quasi-steady state manner. After 10*ms* the deviation to the cable solution is still small (Fig. 2A) because concentrations have changed very little up to this time (Fig. 2C-D), but already around 100*ms* the discrepancy becomes significant due to the fast ions *Na*^+^, *K*^+^ and *Cl*^−^. The *Ca*^2+^ concentration evolves much delayed compared to the other ions because of the much smaller *Ca*^2+^ diffusion coefficient (Fig. 2D). Such a decoupling between ion dynamics can only occur if three or more ions are present, because for two ions the individual dynamics is tightly linked by electroneutrality (ambipolar diffusion). Hence, with more than two ions the concentration dynamics becomes much more complex compared to the case of two ions. The *Na*^+^ influx not only alters the *Na*^+^ concentration (Fig. 2C), but also significantly affects the concentrations of the other ions (Fig. 2C-D). The *K*^+^ concentration is most strongly affected because of the large conductivity fraction *ξ*_*K*_ (see Eq. 12). But also the *Ca*^2+^ concentration at *x* = 1 is reduced by a factor around 2 compared to *x* = 0. The concentration changes for *K*^+^, *Ca*^2+^ and *Cl*^−^ occur due to exchanges with the cell-body, which transiently induce axial currents for these ions (Fig. 2B; only *Na*^+^ and *K*^+^ currents are shown because *Ca*^2+^ and *Cl*^−^ currents are small). At steady state only the *Na*^+^ current survives. At short times the axial *Na*^+^ current becomes shunted because it is used to increase the *Na*^+^ concentration (Fig. 2B).

**Figure 2:**
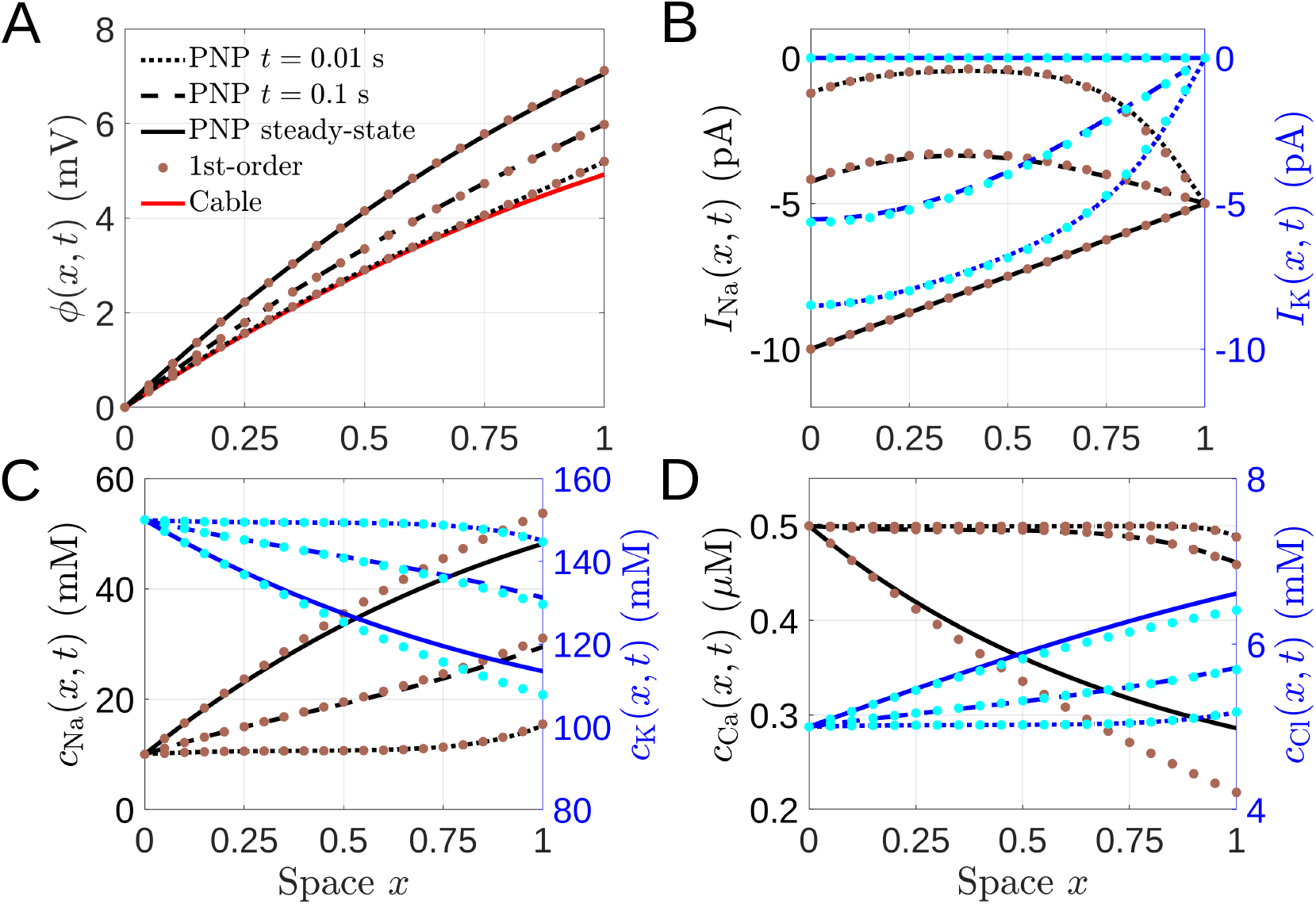
Transient dynamics after the injection of depolarising *Na*^+^ currents. Constant membrane and tip currents *I*_*m,Na*_ = *I*_*tip,Na*_ = −5*pA* are injected for *t >* 0. The PNP results are obtained by numerically solving the PNP equations Eq. 2-3 with COMSOL. First-order approximations indicated by bullets are computed with Eq. 16 and Eq. 15. (A) Voltage dynamics. The red curve is the cable solution in Eq. 17. (B) Axial currents at various times for *Na*^+^ (black, brown) and *K*^+^ (blue, cyan). (C) Concentrations for *Na*^+^ (black, brown) and *K*^+^ (blue, cyan). (D) Concentrations for *Ca*^2+^ (black, brown) and *Cl*^−^ (blue, cyan).

After the transient dynamics we explore steady-state conditions. We first evaluate the steady state with a large depolarising *Na*^+^ membrane current *I*_*m,Na*_ = −50*pA* (Fig. 3). Axial currents at *x* = 1 are zero, *I*_*tip,s*_ = 0. With a larger current compared to (Fig. 2) the discrepancy between first order and PNP increases (Fig. 3 blue vs dashed black curves). Surprisingly, whereas we find large discrepancies for concentrations (the first-order concentration for *Ca*^2+^ becomes even negative) (Fig. 3B-D), there is excellent agreement for the voltage (Fig. 3A). This suggests that concentration errors somehow compensate when contributing to the voltage. We obtain very good agreement with PNP results if voltage and concentrations are computed with the non-linear expressions in Eq. 22, and the first-order method with 10 segments in Eq. 24 (Fig. 3). In contrast, there is significant deviation between PNP and the cable solution (Fig. 3A). The *Na*^+^ influx increases the *Na*^+^ concentration at *x* = 1 by a factor around 10, and also strongly decreases the *K*^+^ concentration (Fig. 3). The *Ca*^2+^ concentration at *x* = 1 becomes reduced by a factor around 5 (Fig. 3C). Voltage and concentration gradients vanish at the tip because axial currents are zero at *x* = 1 (Fig. 3).

**Figure 3:**
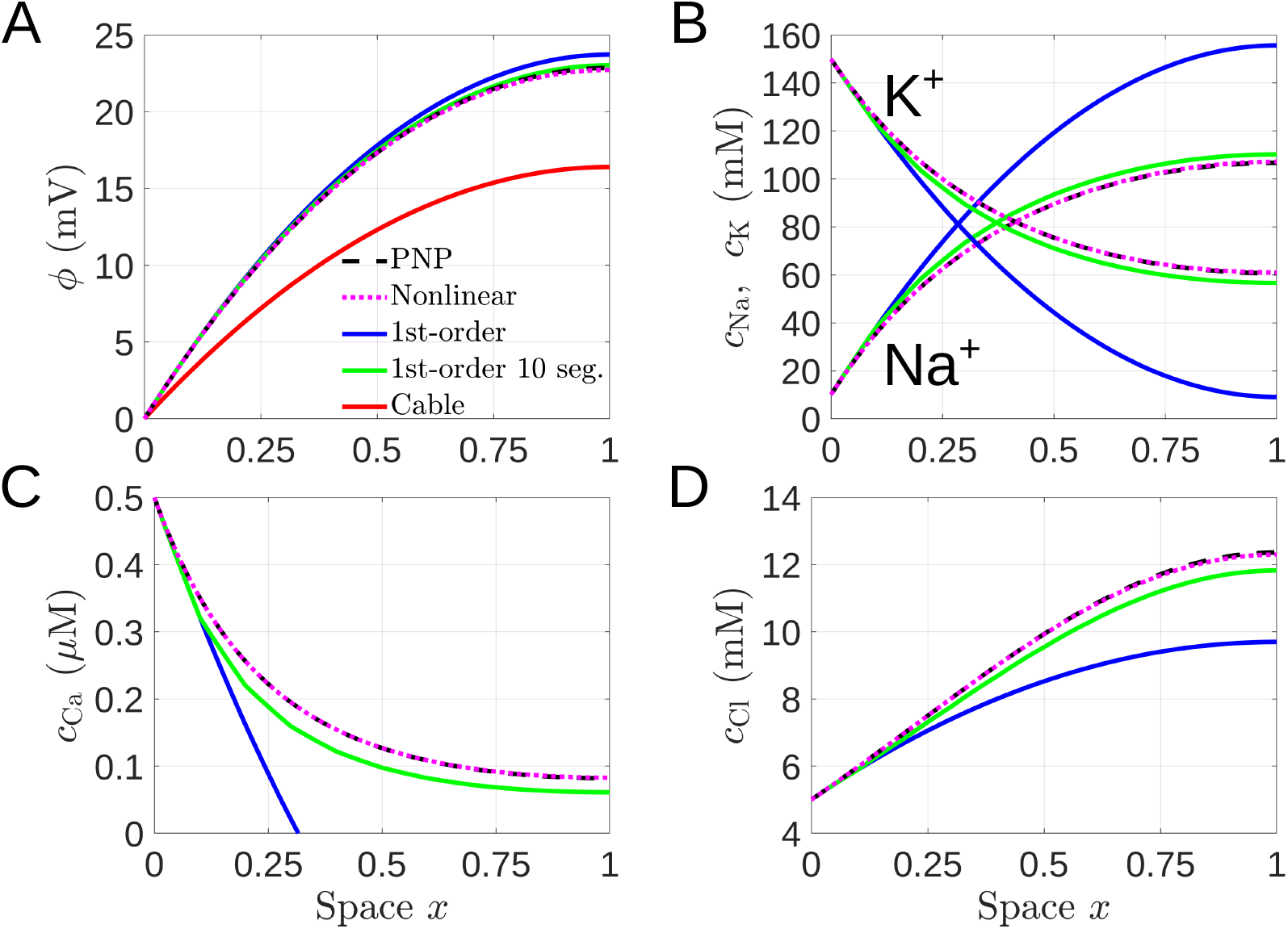
Steady-state results with a large depolarising *Na*^+^ membrane current *I*_*m,Na*_ = −50*pA*. Axial currents at *x* = 1 are zero (*I*_*tip,s*_ = 0). Voltage (A) and concentrations (B-D) are computed with different methods: (1) PNP with COMSOL; (2) non-linear approximations (Eq. 22 and Eq. 23); (3) first-order approximations (Eq. 16 and Eq. 15); (4) first-order and subdividing the cylinder into 10 segments (Eq. 24); (5) cable-equation solution Eq. 17 for the voltage.

Next we explored how the results in Fig. 3 change if depolarisation is achieved by −50*pA* membrane current carried by *Cl*^−^ instead of *Na*^+^. To sustain a large *Cl*− outflux we had to increase the *Cl*− cell-body concentration from 5*mM* to 80*mM*. We maintained the concentrations of positive mobile ions as in Fig. 3, and we adapted the background charge to ensure electroneutrality. The voltage change in Fig. 4A is smaller compared to Fig. 3A due to the larger *Cl*− conductance. Surprisingly, with a *Cl*− current the discrepancy between first-order and PNP results is small for concentrations and large for voltage (Fig. 4), which is opposite to what is observed in Fig. 3. The *Cl*− outflux leads to a more than 10-fold reduction in the *Cl*^−^ concentration at *x* = 1 compared to *x* = 0, which is largely compensated by the reduction of the *K*^+^ concentration (Fig. 4B). The *Na*^+^ concentration at *x* = 1 is reduced by a factor around 2 and *Ca*^2+^ by a factor around 5 (Fig. 4C-D). The osmolarity inside the cylinder is reduced compared to Fig. 3 with a *Na*^+^ current. Thus, a depolarising *Cl*^−^ current is beneficial to prevent a large increase in osmotic pressure.

**Figure 4:**
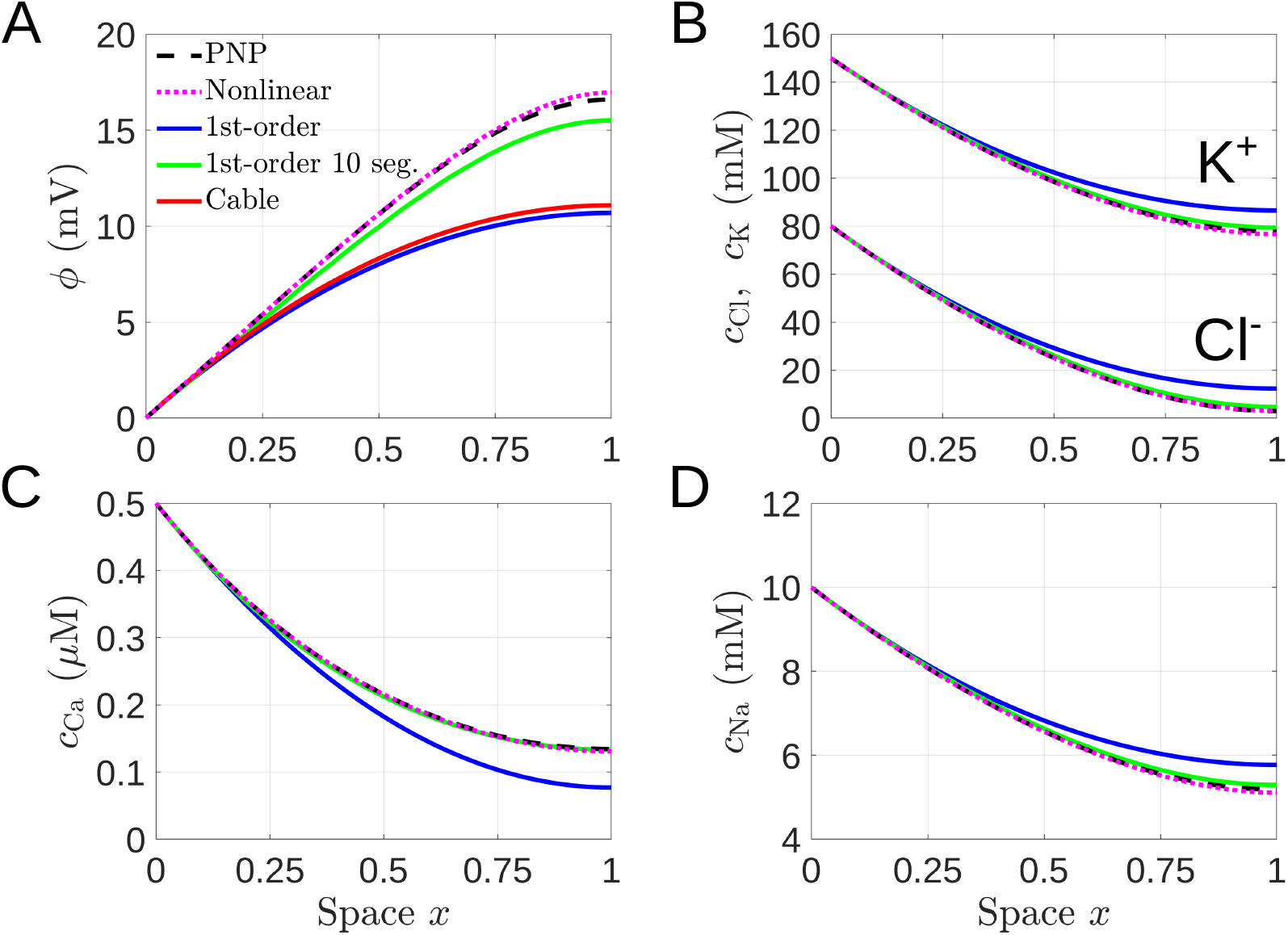
Steady-state results with a large depolarizing *Cl*^−^ membrane current *I*_*m,Cl*_ = −50*pA*. Axial currents at *x* = 1 are zero. Voltage (A) and concentrations (B-D) are computed as described in Fig. 3. To sustain a large *Cl*^−^ outflux we increased the cell-body *Cl*^−^ concentration from 5*mM* in Fig. 3 to 80*mM*. Cell-body concentrations of *Na*^+^, *K*^+^ and *Ca*^2+^ did not change. We adapted the immobile background charge to ensure electroneutrality.

Finally, we investigated more complex scenarios with multiple ionic currents (Fig. 5). We show two examples, one where the cable equation completely fails, and one where it is correct. In the first example we chose currents such that the total current *I*(*x*) vanishes. In this case the cable equation erroneously predicts zero voltage change, contrary to PNP and analytic results (Fig. 5)A). In the second example we chose the individual currents distributed according to Eq. 25 such that concentrations remain constant (not shown). After switching on the currents, PNP and first-order solution jump to the cable result as in Fig. 2, but now they do not evolve any further because concentrations remain constant (Fig. 5)B).

**Figure 5:**
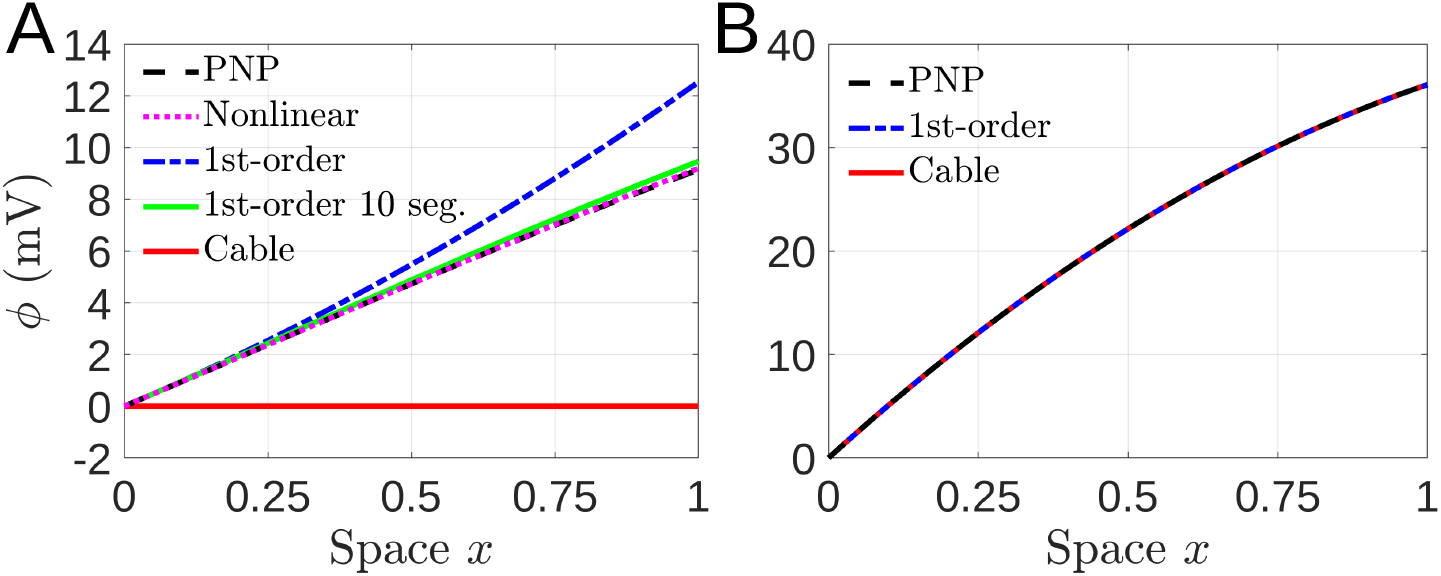
Steady-state voltage with multiple ionic currents. Voltages are computed as described in Fig. 3. (A) Zero total current *I*(*x*) = 0 with *I*_*tip,Na*_ = −50*pA, I*_*tip,K*_ = 30*pA, I*_*tip,Cl*_ = 20*pA* and *I*_*m,Na*_ = 20*pA, I*_*m,K*_ = −10*pA, I*_*m,Cl*_ = −10*pA*. (B) Individual currents are distributed according to Eq. 25 with total currents *I*_*m*_ = −50*pA* and *I*_*tip*_ = −30*pA*. With this choice concentrations remain constant (not shown).

## 6 Conclusion

In this work we used the electroneutrality approximation to reduce the PNP model to a system of cross-diffusion equations for ionic concentrations. These equations expose how current and concentration gradient of one ion species affects the other species. We derived analytic result that explicitly show how axial and membrane currents alter voltage and ion concentrations in a thin cylindrical domain.

After the current onset, the voltage changes in a biphasic manner: it first jumps to a profile that is given by the steady-state cable solution with constant initial concentrations, and then co-evolves in a quasi-steady state manner with the diffusion controlled ion concentrations (Fig. 2). Such a biphasic voltage behaviour has also been observed numerically for a more complex PNP model after the opening of ion channels in a small cardiac dyad [30]. With only two ions the individual concentrations are tightly coupled due to electroneutrality (ambipolar diffusion). In contrast, with three or more ions different combinations of ion concentrations ensure electroneutrality, and the ion dynamics becomes much more rich and complex (Fig. 3-Fig. 4). For example, a *Na*^+^ membrane current not only changes the *Na*^+^ concentration, as in most electrodiffusion models, but also strongly alters the concentrations of *K*^+^, *Ca*^2+^ and *Cl*^−^ (Fig. 3). This is particularly relevant for *Ca*^2+^ because *Ca*^2+^ is an ubiquitous regulator of cellular functions, and many processes are extremely sensitive to changes in the *Ca*^2+^ concentration [50]. Hence, we conclude that cellular functions might be affected even if no *Ca*^2+^ channels are present.

We derive time-dependent formula for voltage and concentration profiles in the limit of small concentration changes (Eq. 16, Eq. 15), and higher-order expressions for steady-state voltage and concentrations (Eq. 22, Eq. 23). Hence, the first-order results provide corrections to the cable model and the Rall theory that assume constant concentrations [41, 40]. We show that the first-order results can be used to obtain accurate predictions of steady-state voltage and concentrations by subdividing the cylinder into small segments. This recursive method does not rely on solving PNP equations, and could be implemented into the popular simulation software NEURON [43] to improve the modelling of neuronal or dendritic domains.

Cross-diffusion theory has been used in the context of pattern formation with reaction-diffusion systems, multicomponent ion diffusion in silicates or non-equilibrium thermodynamics [51, 52, 53]. Our analysis might open new avenues where this theory is applied in more detail to neuroscience. Future research will have to develop a feedback mechanism where the injected currents are not constant but are modified by voltage and concentration changes. Another future challenge will be to generalize the multi-segment recursive framework from steady-state to transient dynamics. Whereas the actual cross-diffusion model describes coarse-grained voltage and concentration changes, it does not allow for Debye layer analysis, nor does it accurately describe fast voltage dynamics that depends on the membrane capacitance, such as action potentials. To study the Debye layer one has to engage the PNP model, and for action potentials classical models with constant concentrations are often sufficiently accurate.

## 7 APPENDIX

### 7.1 Eigenvectors and eigenvalues of the matrix Λ

A set of unnormalized eigenvectors 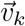 of **Λ**_*si*_ = *ν*_*s*_*δ*_*si*_ − *ξ*_*s*_*ν*_*i*_ to eigenvalues *λ*_*k*_ (we choose *λ*_*N*_ = 0) are

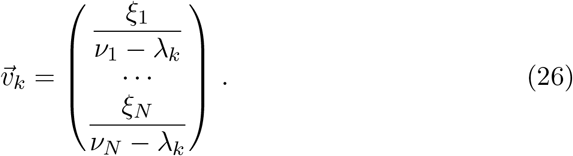

The non-zero eigenvalue for *N* = 2 ions is

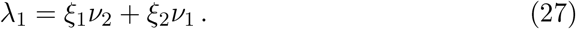

The non-zero eigenvalues for *N* = 3 ions are

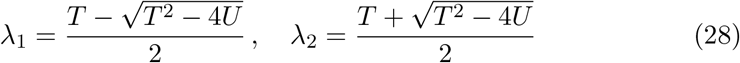

with *T* = ∑_*s*_ *ν*_*s*_(1 − *ξ*_*s*_) and *U* = *ξ*_1_*ν*_2_*ν*_3_ + *ξ*_2_*ν*_1_*ν*_3_ + *ξ*_3_*ν*_1_*ν*_2_.

### 7.2 Steady-state polynomial expansion of *γ*(*x*)

We derive the steady-state polynomial expansion of *γ*_*s*_(*x*) = *z*_*s*_*ρ*_*s*_(*x*) around *x* = 0 as a function of the steady-state currents *I*_*s*_(*x*). To simplify the notation we introduce 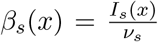 and *β*(*x*) = Σ_*s*_*β*_*s*_(*x*). By combining Eq. 21 and Eq. 1 we find that *γ*_*s*_(*x*) satisfy

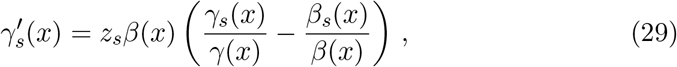

where *γ*(*x*) = ∑ _*s*_ *γ*_*s*_(*x*). With Eq. 29 we obtain for the coefficients of the polynomial approximation 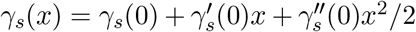 the recursive expressions (*γ*_*s*_(0) are given by the Dirichlet boundary values)

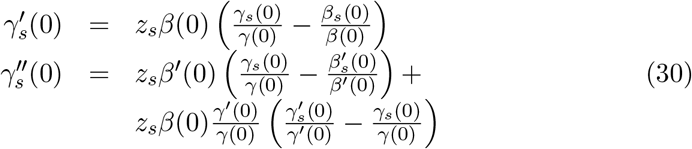

The background charge *ρ*_*b*_ affects Eq. 30 through the values *γ*_*s*_(0). This procedure can be extended to compute higher-order polynomial approximations.

Note that the coefficients in Eq. 30 are exact and not a first-order approximation. An alternative method is to compute 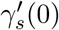 and 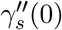 using the first-order result in Eq. 16. This gives identical values for 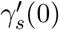 because Eq. 6 at *x* = 0 coincides with the first-order approximation. In contrast, the values for 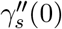 will differ because the derivative of Eq. 6 at *x* = 0 involves 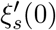, which is zero for the first-order approximation. For example, with *I*_*m,s*_ = 0 we obtain 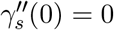 with Eq. 16 and 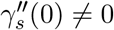 with Eq. 30.

## References

[1] G. A. Silva, “Neuroscience nanotechnology: progress, opportunities and challenges,” Nature reviews neuroscience, vol. 7, no. 1, pp. 65–74, 2006.

[2] D. Holcman and R. Yuste, “The new nanophysiology: regulation of ionic flow in neuronal subcompartments,” Nature Reviews Neuroscience, vol. 16, no. 11, pp. 685–692, 2015.

[3] G. Marmont, “Studies on the axon membrane. i. a new method,” Journal of cellular and comparative physiology, vol. 34, no. 3, pp. 351–382, 1949.

[4] J. W. Moore and K. S. Cole, “Voltage clamp techniques,” Physical techniques in biological research, vol. 6, pp. 263–321, 1963.

[5] A. L. Hodgkin and A. F. Huxley, “A quantitative description of membrane current and its application to conduction and excitation in nerve,” The Journal of physiology, vol. 117, no. 4, p. 500, 1952.

[6] H. H. Yang and F. St-Pierre, “Genetically encoded voltage indicators: opportunities and challenges,” Journal of Neuroscience, vol. 36, no. 39, pp. 9977–9989, 2016.

[7] Y. Xu, P. Zou, and A. E. Cohen, “Voltage imaging with genetically encoded indicators,” Current opinion in chemical biology, vol. 39, pp. 1–10, 2017.

[8] K. Jayant, J. J. Hirtz, I. J.-L. Plante, D. M. Tsai, W. D. A. M. De Boer, A. Semonche, D. S. Peterka, J. S. Owen, O. Sahin, K. L. Shepard, and R. Yuste, “Targeted intracellular voltage recordings from dendritic spines using quantum-dot-coated nanopipettes,” Nature Nanotech., vol. 12, pp. 335–342, 2017.

[9] J. Mc Hugh, S. Makarchuk, D. Mozheiko, A. Fernandez-Villegas, G. S. Kaminski Schierle, C. F. Kaminski, U. F. Keyser, D. Holcman, and N. Rouach, “Diversity of dynamic voltage patterns in neuronal dendrites revealed by nanopipette electrophysiology,” Nanoscale, vol. 15, pp. 12245–12254, 2023.

[10] G. G. Somjen, Ions in the brain: normal function, seizures, and stroke. Oxford University Press, 2004.

[11] B. Hille, Ionic Channels of Excitable Membranes. Sinauer, Sunderland, third ed., 2001.

[12] R. M. Paredes, J. C. Etzler, L. T. Watts, W. Zheng, and J. D. Lechleiter, “Chemical calcium indicators,” Methods, vol. 46, no. 3, pp. 143–151, 2008.

[13] C. Grienberger and A. Konnerth, “Imaging calcium in neurons,” Neuron, vol. 73, no. 5, pp. 862–885, 2012.

[14] S. Kirischuk, V. Parpura, and A. Verkhratsky, “Sodium dynamics: another key to astroglial excitability?,” Trends in neurosciences, vol. 35, no. 8, pp. 497–506, 2012.

[15] N. J. Gerkau, R. Lerchundi, J. S. Nelson, M. Lantermann, J. Meyer, J. Hirrlinger, and C. R. Rose, “Relation between activity-induced intracellular sodium transients and atp dynamics in mouse hippocampal neurons,” The Journal of physiology, vol. 597, no. 23, pp. 5687–5705, 2019.

[16] G. Ullah, J. R. Cressman Jr, E. Barreto, and S. J. Schiff, “The influence of sodium and potassium dynamics on excitability, seizures, and the stability of persistent states: Ii. network and glial dynamics,” Journal of computational neuroscience, vol. 26, no. 2, pp. 171–183, 2009.

[17] J. R. Cressman Jr, G. Ullah, J. Ziburkus, S. J. Schiff, and E. Barreto, “The influence of sodium and potassium dynamics on excitability, seizures, and the stability of persistent states: I. single neuron dynamics,” Journal of computational neuroscience, vol. 26, no. 2, pp. 159–170, 2009.

[18] Z. Schuss, B. Nadler, and R. S. Eisenberg, “Derivation of poisson and nernst-planck equations in a bath and channel from a molecular model,” Phys. Rev. E, vol. 64, p. 036116, Aug 2001.

[19] B. Eisenberg, “Ionic channels in biological membranes-electrostatic analysis of a natural nanotube,” Contemporary Physics, vol. 39, no. 6, pp. 447–466, 1998.

[20] D. Chen and R. Eisenberg, “Charges, currents, and potentials in ionic channels of one conformation,” Biophysical journal, vol. 64, no. 5, pp. 1405–1421, 1993.

[21] T. M. Kamsma, W. Q. Boon, T. ter Rele, C. Spitoni, and R. van Roij, “Iontronic neuromorphic signaling with conical microfluidic memristors,” Phys. Rev. Lett., vol. 130, p. 268401, Jun 2023.

[22] H. Row, J. B. Fernandes, K. K. Mandadapu, and K. Shekhar, “Spatiotemporal dynamics of ionic reorganization near biological membrane interfaces,” Physical Review Research, vol. 7, no. 1, p. 013185, 2025.

[23] D. Ben-Yaakov, D. Andelman, D. Harries, and R. Podgornik, “Beyond standard poisson–boltzmann theory: ion-specific interactions in aqueous solutions,” J. Phys.: Condens. Matter, vol. 21, p. 424106, sep 2009.

[24] T. Markovich, D. Andelman, and R. Podgornik, Charged Membranes: Poisson–Boltzmann Theory, The DLVO Paradigm, and Beyond, pp. 99–128. CRC Press, 1st ed., 2021.

[25] J. Pods, J. Schönke, and P. Bastian, “Electrodiffusion models of neurons and extracellular space using the poisson-nernst-planck equations—numerical simulation of the intra- and extracellular potential for an axon model,” Biophysical Journal, vol. 105, no. 1, pp. 242–254, 2013.

[26] C. L. Lopreore, T. M. Bartol, J. S. Coggan, D. X. Keller, G. E. Sosinsky, M. H. Ellisman, and T. J. Sejnowski, “Computational modeling of three-dimensional electrodiffusion in biological systems: application to the node of ranvier,” Biophysical journal, vol. 95, no. 6, pp. 2624–2635, 2008.

[27] D. H. Song, J. Park, L. L. Maurer, W. Lu, M. A. Philbert, and A. M. Sastry, “Biophysical significance of the inner mitochondrial membrane structure on the electrochemical potential of mitochondria,” Physical Review E, vol. 88, no. 6, p. 062723, 2013.

[28] T. Lagache, K. Jayant, and R. Yuste, “Electrodiffusion models of synaptic potentials in dendritic spines,” J Comput Neurosci, vol. 47, pp. 77–89, 2019.

[29] F. Boahen and N. Doyon, “Modelling dendritic spines with the finite element method, investigating the impact of geometry on electric and calcic responses,” J. Math. Biol., vol. 81, no. 2, pp. 517–547, 2020.

[30] K. Horgmo Jæ ger and A. Tveito, “Electrodiffusion dynamics in the cardiomyocyte dyad at nano-scale resolution using the poisson-nernst-planck (pnp) equations,” PLOS Computational Biology, vol. 21, no. 6, p. e1013149, 2025.

[31] C. Guerrier, T. D. Toth, N. Galtier, and K. Haas, “An algorithm based on a cable-nernst planck model predicting synaptic activity throughout the dendritic arbor with micron specificity,” Neuroinformatics, vol. 21, no. 1, pp. 207–220, 2023.

[32] J. J. Jasielec, “Electrodiffusion phenomena in neuroscience and the nernst–planck–poisson equations,” Electrochem, vol. 2, no. 2, pp. 197–215, 2021.

[33] N. Qian and T. Sejnowski, “An electro-diffusion model for computing membrane-potentials and ionic concentrations in branching dendrites, spines and axons.,” Biological Cybernetics, vol. 62, no. 1, pp. 1–15, 1989.

[34] M. Breit and G. Queisser, “The necessary modeling detail for neuronal signaling: Poisson–nernst–planck and cable equation models in one and three dimensions,” SIAM Journal on Applied Mathematics, vol. 81, no. 2, pp. 530–550, 2021.

[35] G. Halnes, I. Ostby, K. Pettersen, S. Omholt, and G. Einevoll, “Electrodiffusive model for astrocytic and neuronal ion concentration dynamics.,” PLoS Comput Biol, vol. 9, no. 12, p. e1003386, 2013.

[36] Y. Mori, “A multidomain model for ionic electrodiffusion and osmosis with an application to cortical spreading depression,” Physica D: Nonlinear Phenomena, vol. 308, pp. 94–108, 2015.

[37] M. J. Sætra and Y. Mori, “An electrodiffusive network model with multicompartmental neurons and synaptic connections,” PLOS Computational Biology, vol. 20, no. 11, p. e1012114, 2024.

[38] J. Reisert and J. Reingruber, “Ca2+-activated cl-current ensures robust and reliable signal amplification in vertebrate olfactory receptor neurons.,” Proc Natl Acad Sci U S A, vol. 116, no. 3, pp. 1053–1058, 2019.

[39] I. Segev, J. Rinzel, G. M. Shepherd, and A. Borst, “The theoretical foundation of dendritic function,” Trends in Neurosciences, vol. 18, no. 11, pp. 512–512, 1995.

[40] W. Rall et al., “Core conductor theory and cable properties of neurons,” Chapter in Handbook of physiology, vol. 1, no. part 1, pp. 39–98, 1977.

[41] W. Rall, “Theory of physiological properties of dendrites,” Annals of the New York Academy of Sciences, vol. 96, no. 4, pp. 1071–1092, 1962.

[42] J. Jack, D. Noble, and R. Tsien, Electrical current flow in excitable cells. Oxford University Press, 1975.

[43] M. L. Hines and N. T. Carnevale, “The neuron simulation environment,” Neural Computation, vol. 9, pp. 1179–1209, 08 1997.

[44] L. P. Savtchenko, L. Bard, T. P. Jensen, J. P. Reynolds, I. Kraev, N. Medvedev, M. G. Stewart, C. Henneberger, and D. A. Rusakov, “Disentangling astroglial physiology with a realistic cell model in silico,” Nature Communications, vol. 9, no. 1, 2018.

[45] J. Keener and J. Sneyd, Mathematical Physiology. Springer-Verlag New York Inc., 1998.

[46] Y. Mori, G. I. Fishman, and C. S. Peskin, “Ephaptic conduction in a cardiac strand model with 3d electrodiffusion,” Proceedings of the National Academy of Sciences, vol. 105, no. 17, pp. 6463–6468, 2008.

[47] G. S. Hartley and C. Robinson, “The diffusion of colloidal electrolytes and other charged colloids,” Proceedings of the Royal Society of London. Series A, Containing Papers of a Mathematical and Physical Character, vol. 134, pp. 20–35, 11 1931.

[48] L. J. Gosting, “Measurement and interpretation of diffusion coefficients of proteins,” vol. 11 of Advances in Protein Chemistry, pp. 429–554, Academic Press, 1956.

[49] M. A. Lieberman and A. J. Lichtenberg, Principles of Plasma Discharges and Materials Processing. John Wiley and Sons, Ltd, second ed., 2005.

[50] M. J. Berridge, “Neuronal calcium signaling,” Neuron, vol. 21, no. 1, pp. 13–26, 1998.

[51] V. K. Vanag and I. R. Epstein, “Cross-diffusion and pattern formation in reaction–diffusion systems,” Phys. Chem. Chem. Phys., vol. 11, pp. 897–912, 2009.

[52] A. C. Lasaga, “Multicomponent exchange and diffusion in silicates,” Geochimica et Cosmochimica Acta, vol. 43, no. 4, pp. 455–469, 1979.

[53] S. R. De Groot and P. Mazur, Non-equilibrium thermodynamics. Courier Corporation, 2013.

